# Semiparametric methods for estimation of a non-linear exposure–outcome relationship using instrumental variables with application to Mendelian randomization

**DOI:** 10.1101/103986

**Authors:** James R Staley, Stephen Burgess

**Affiliations:** Cardiovascular Epidemiology Unit, Department of Public Health and Primary Care, University of Cambridge, United Kingdom

## Abstract

Mendelian randomization, the use of genetic variants as instrumental variables (IV), can test for and estimate the causal effect of an exposure on an outcome. Most IV methods assume that the function relating the exposure to the expected value of the outcome (the exposure-outcome relationship) is linear. However, in practice this assumption may not hold. Indeed, often the primary question of interest is to assess the shape of this relationship. We present two novel IV methods for investigating the shape of the exposure-outcome relationship: a fractional polynomial method and a piecewise linear method. We divide the population into strata using the exposure distribution, and estimate a causal effect, referred to as a localized average causal effect (LACE), in each stratum of population. The fractional polynomial method performs meta-regression on these LACE estimates. The piecewise linear method estimates a continuous piecewise linear function, the gradient of which is the LACE estimate in each stratum. Both methods were demonstrated in a simulation study to estimate the true exposure-outcome relationship well, particularly when the relationship was a fractional polynomial (for the fractional polynomial method) or was piecewise linear (for the piecewise linear method). The methods were used to investigate the shape of relationship of body mass index with systolic blood pressure and diastolic blood pressure.

Availability and implementation: https://github.com/jrs95/nlmr

## Introduction

Often the shape of association between an exposure and an outcome is non-linear. For example, the observed association between body mass index (BMI) and all-cause mortality in a Western context is J-shaped (or U-shaped), as risk of mortality is increased for individuals at both ends of the BMI distribution [Flegal et al., 2013]. However, particularly for underweight individuals, this could reflect either reverse causality or confounding, rather than a true causal effect of low BMI increasing mortality risk. Instrumental variable (IV) methods can be used to distinguish between correlation and causation. However, these methods typically assume that the exposure-outcome relationship is linear when estimating a causal effect [Hernán and Robins, 2006]. In many cases, investigating the shape of the exposure-outcome relationship is the primary aim of a study. This can be used to define treatment thresholds for pharmaceutical interventions or health guidelines.

A natural way of tackling the non-linearity problem in IV analysis is to perform a two-stage analysis similar to the well-known two-stage least squares method, except fitting a non-linear function in the second stage [Horowitz, 2011; Newey and Powell, 2003]. However, this approach requires the instrument and any covariates included in the first-stage model to explain a large proportion of variance in the exposure, as information for assessing the shape of relationship between the exposure and outcome will only be available for the fitted values of the exposure from the first-stage regression. If the proportion of variance in the exposure explained by the IV is small then observing non-linearity for this limited range of values is unlikely. In Mendelian randomization, the use of genetic variants as instrumental variables, genetic variants typically only explain a small percentage of the variance in the exposure (usually in the region of 1–4%) [Burgess and Thompson, 2015; Ebrahim and Smith, 2008].

Two approaches for addressing non-linearity in the context of Mendelian randomization have recently been proposed [Burgess et al., 2014; Silverwood et al., 2014]. Burgess et al. assessed the consequences of performing a linear IV analysis when the exposure-outcome relationship truly was non-linear, as well as stratifying individuals using the exposure distribution to obtain IV estimates, referred to as localized average causal effects (LACE), in each stratum. Silverwood et al. performed meta-regression of LACE estimates across strata to examine whether a quadratic rather than a linear model was a better fit for relationships between alcohol consumption and a variety of cardiovascular markers.

In this paper, we present two novel semiparametric methods for investigating the shape of the exposure-outcome relationship using instrumental variable analysis developed for use in Mendelian randomization. The first is based on fractional polynomials [Royston and Altman, 1994; Royston et al., 1999], whereas the second fits a piecewise linear function. We also propose a test for non-linearity based on the fractional polynomial method, and assess the impact of varying the number of strata of the exposure distribution used to test for nonlinearity and to estimate non-linear relationships. We illustrate the methods using data from UK Biobank [Sudlow et al., 2015], a large UK-based cohort, to investigate the shape of the relationship between BMI and blood pressure using Mendelian randomization.

## Methods

### Stratifying on the IV-free exposure

We define the exposure-outcome relationship as the function relating the exposure to the expected value of the outcome. We initially assume that this function is homogeneous for all individuals in the population, and return to its interpretation in case of heterogeneity in the discussion.

To assess the shape of association between exposure *X* and outcome *Y* using a single instrument *G*, we first stratify the population using the exposure distribution. If we were to stratify on the exposure directly, then an association between the IV and outcome might be induced even if it were not present in the original data, thus invalidating the IV assumptions [Didelez and Sheehan, 2007]. This problem can be avoided by instead stratifying on the residual variation in the exposure after conditioning on the IV, assuming that the effect of the IV on the exposure is linear and constant for all individuals across the entire of the exposure distribution [Burgess et al., 2014]. In econometrics, this residual is known as a control function [Arellano, 2003]. We calculate this residual by performing linear regression of the exposure on the IV, and then setting the value of the IV to 0. We refer to this as the IV-free exposure. It is the expected value of the exposure that would be observed if the individual had an IV value of 0, and can be interpreted as the non-genetic component of the exposure.

In each stratum of the IV-free exposure, we estimate the LACE as a ratio of coefficients: the IV association with the outcome divided by the IV association with the exposure. The assumption that the effect of the IV on the exposure is constant is a stronger version of the monotonicity assumption [Angrist et al., 1996], and hence the LACE are local average treatment effects (also called complier-averaged causal effects [Yau and Little, 2001]) for each stratum [Imbens and Angrist, 1994]. We then proceed to estimate the exposure-outcome relationship from these LACE estimates using two approaches: the first based on fractional polynomials, and the second a piecewise linear function.

### Fractional polynomial method

The fractional polynomial method consists of meta-regression of the LACE estimates against the mean of the exposure in each stratum in a flexible semiparametric framework [Bagnardi et al., 2004; Thompson and Sharp, 1999]. Fractional polynomials are a family of functions that can be used to fit complex relationships for a single covariate [Royston and Altman, 1994]. The standard powers used when modelling using fractional polynomials are *P* = {−2, −1, −0.5, 0, 0.5, 1, 2, 3}, where the power of 0 refers to the (natural) log function. These powers are used throughout this paper. Fractional polynomials of degree 1 are defined as:

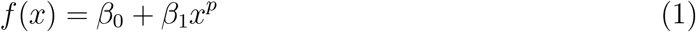

where *p ∈ P*. Similarly, fractional polynomials of degree 2 are defined as:

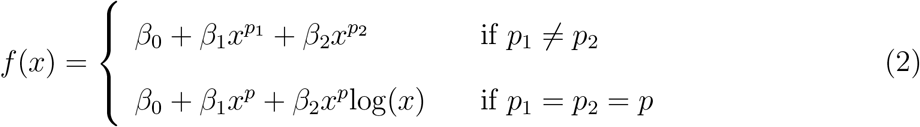

where *p*_1_, *p*_2_ ∈ *P*. In both cases, *x*^0^ is interpreted as log(*x*). As fractional polynomials of degree larger than 2 are rarely required in practice, these were not considered in this paper [Royston and Altman, 1994]. Since a causal effect is an estimate of the derivative of the exposure-outcome relationship [Small, 2014], we fit the LACE estimates using the derivative of the fractional polynomial function (from either (1) or (2)).

The method proceeds as follows. First, we calculate the IV-free exposure, and stratify the population based on quantiles of its distribution. Secondly, the LACE estimate is calculated in each stratum as a ratio of coefficients (the LACE estimate for stratum *k* is 
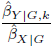
, where 
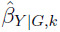
, is the estimated association of the IV with the outcome in stratum *k* and 
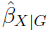
 is the estimated association of the IV with the exposure in the whole population), and the standard error of the LACE estimate is computed as 
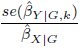

(the first term of the delta method approximation [Thomas et al., 2007]). Third, these LACE estimates are meta-regressed against the mean of the exposure in each stratum using the derivative of the fractional polynomial function as the model relating the LACE estimates to the exposure values. The original fractional polynomial function then represents the exposure-outcome relationship. As this function is constructed from the LACE estimates, the intercept of the exposure-outcome curve cannot be estimated and must be set arbitrarily. If it is set to zero at a reference value (for instance, the mean of the exposure distribution), then the value of the function represents the expected difference in the outcome compared with this reference value when the exposure is set to different values.

Confidence intervals for the exposure-outcome curve can be computed arithmetically under a normal assumption either using the estimated standard errors from the meta-regression or by bootstrapping the second and third steps from above (we maintain the strata and the-estimate of the IV on the exposure as in the original data, and estimate the associations of the IV with the outcome in bootstrapped samples for each stratum).

To explore a range of possible parametric forms, we fit all possible fractional polynomial models of degrees 1 and 2, and select the best-fitting one based on the likelihood. A fractional polynomial of degree 2 is preferred over one of degree 1 if the twice the difference in the log-likelihood is greater than the 95th percentile point of a 
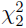
 distribution for the best-fitting fractional polynomial in each class [Royston and Altman, 1994].

### Piecewise linear method

Another way of estimating the exposure-outcome relationship is to use a piecewise linear approach. The exposure-outcome relationship is estimated as piecewise linear function with each stratum contributing a line segment whose gradient is the LACE estimate for that stratum. The function is constrained to be continuous, so that each line segment begins where the previous segment finished. As in the fractional polynomial method, while the intercept for each line segment is fixed by the previous line segment, the overall intercept of the exposure-outcome curve cannot be estimated and must be set arbitrarily.

Confidence intervals are estimated by bootstrapping the IV associations with the outcome as in the fractional polynomial approach. For a 95% confidence interval, the piecewise linear method is performed for each bootstrapped dataset, and then the 2.5th and 97.5th percentiles of the function are taken at selected points across the exposure distribution; we chose the mean exposure values in each of the strata.

### Tests of non-linearity

There are already two proposed methods in this framework for testing whether a non-linear exposure-outcome model fits the data better than a linear model. The first is a heterogeneity test using Cochran’s Q statistic to assess whether the LACE estimates differ more than would be expected by chance. The second is a trend test where the LACE estimates are meta-regressed against the mean value of the exposure in each strata; this is equivalent to fitting a quadratic exposure-outcome model. A more flexible version of this method is to test the best-fitting fractional polynomial of degree 1 against the linear model. This can be achieved by comparing twice the difference in the log-likelihood between the linear model and the best fitting fractional polynomial of degree 1 with a 
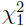
 distribution.

These methods are available in the nlmr R package (https://github.com/jrs95/nlmr).

### Simulation study

To assess the performance of these methods in realistic scenarios for Mendelian randomization, we performed a simulation study. We simulated data for 10,000 individuals for an IV *G*, a continuous exposure *X* that takes only positive values, a continuous outcome *Y*, and a confounder *U* (assumed to be unmeasured). The data generating model for individual *i* is:

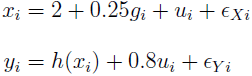

where *g_i_* ~ Bin(2, 0.3), *ϵ _Xi_*~ Exp(1), *u_i_* ~ Unif(0,1), *ϵ_Yi_* ~ *N*(0, 1), and *h*(*x_i_*) is the function relating the exposure to the outcome (the exposure-outcome relationship). Exposure values were taken to be positive and away from zero so that the outcome takes sensible values for log and negative power functions.

### Choice of exposure-outcome model

For the fractional polynomial method, all possible fractional polynomials of degrees 1 and 2 were considered as the functional form of the exposure-outcome relationship. Combinations of effect sizes for the *β* parameters were chosen ranging from 0 to 2. For fractional polynomials of degree 2, we also considered effects in opposing directions for *β*_1_ and *β*_2_; these simulations yielded similar results to those discussed here (results not shown). Fixed-effects meta-regression was used in the simulations, however, random-effects meta-regression yielded similar results (results not shown).

For the piecewise linear method and for comparisons between methods, linear, quadratic, square root, and logarithm functions were considered as the functional form of the exposure-outcome relationship, as well as a threshold model:

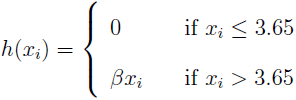

### Evaluating the performance of the methods

To evaluate the fractional polynomial method, we first fitted the correct fractional polynomial model (that is, with the correct degree and powers) and assessed the bias and coverage of the effect parameter estimates. Subsequently, we fitted all fractional polynomials of the same degree and selected the best-fitting polynomial based on the likelihood. We assessed the proportion of simulations where the best-fitting fractional polynomial was the correct fractional polynomial. If the correct fractional polynomial was not the best-fitting fractional polynomial, we tested whether it was in the group of fractional polynomials that fit the data almost as well as the best-fitting polynomial; defined as those fractional polynomials where twice the difference in the log-likelihood (compared with the best-fitting polynomial) was less than the 90th percentile point of a 
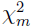
 distribution, where *m* = 1 for comparing fractional polynomials of degree 1 and *m* = 2 for comparing polynomials of degree 2.

To evaluate the piecewise linear method, we first compared the outcome estimates at the mean exposure value in each quantile to the values of the true model at the same points. The coverages of the bootstrapped 95% confidence intervals were also evaluated at these points.

For comparing the fit of the fractional polynomial and piecewise linear models, we used the following heuristic function:

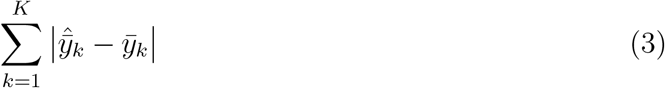

where summation is across the *K* quantile groups, and 
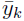
 is the expected value of the outcome evaluated at the mean value of the exposure in each quantile group.

### Varying the number of strata

In the initial simulations, the population was split based on the IV-free exposure into decile groups. Further simulations were performed varying the number of strata using 5, 10, 50 and 100 quantile groups. Tests of non-linearity were performed to assess the impact of the number of strata on the empirical power of each test. The empirical power of each test was reported as the proportion of simulation replicates with *p*-value less than 0.05. The heuristic function (3) was calculated based on 10 deciles for each number of strata.

For each simulation and set of parameters, 500 replications were performed. Bootstrap 95% confidence intervals were generated using 500 bootstrap samples. All analyses were performed using R version 3.0.2.

### Additional simulations to assess impact of violations of assumptions

We performed additional simulations in which the underlying assumptions that the effect of the IV on the exposure and the effect of the exposure on the outcome are fixed and independent were relaxed. In these simulations, we assessed both modelling assumptions by allowing the effect of the IV on the exposure to vary (by drawing the effect parameter from a normal distribution *N*(0.25, 0.1^2^) for each individual in the population), and allowing the exposure-outcome relationship to vary (by drawing the causal parameter from a normal distribution *N*(*β*, 0.2^2^) for each individual in the population). We assessed the impact of allowing each of these parameters to vary separately and both to vary together. In addition, we also allowed variation in both parameters to be correlated by drawing the parameters from a bivariate normal distribution with correlation 0.2. For fractional polynomials of degree 2, only the causal parameter for the second polynomial was allowed to vary across individuals.

## Results

### Fractional polynomial method

Comparisons of fractional polynomials for all powers are provided in Supplementary Table S1 (degree 1) and Supplementary Table S2 (degree 2); a summary of results for the most commonly encountered powers is given in Table 1.

**Table 1.**
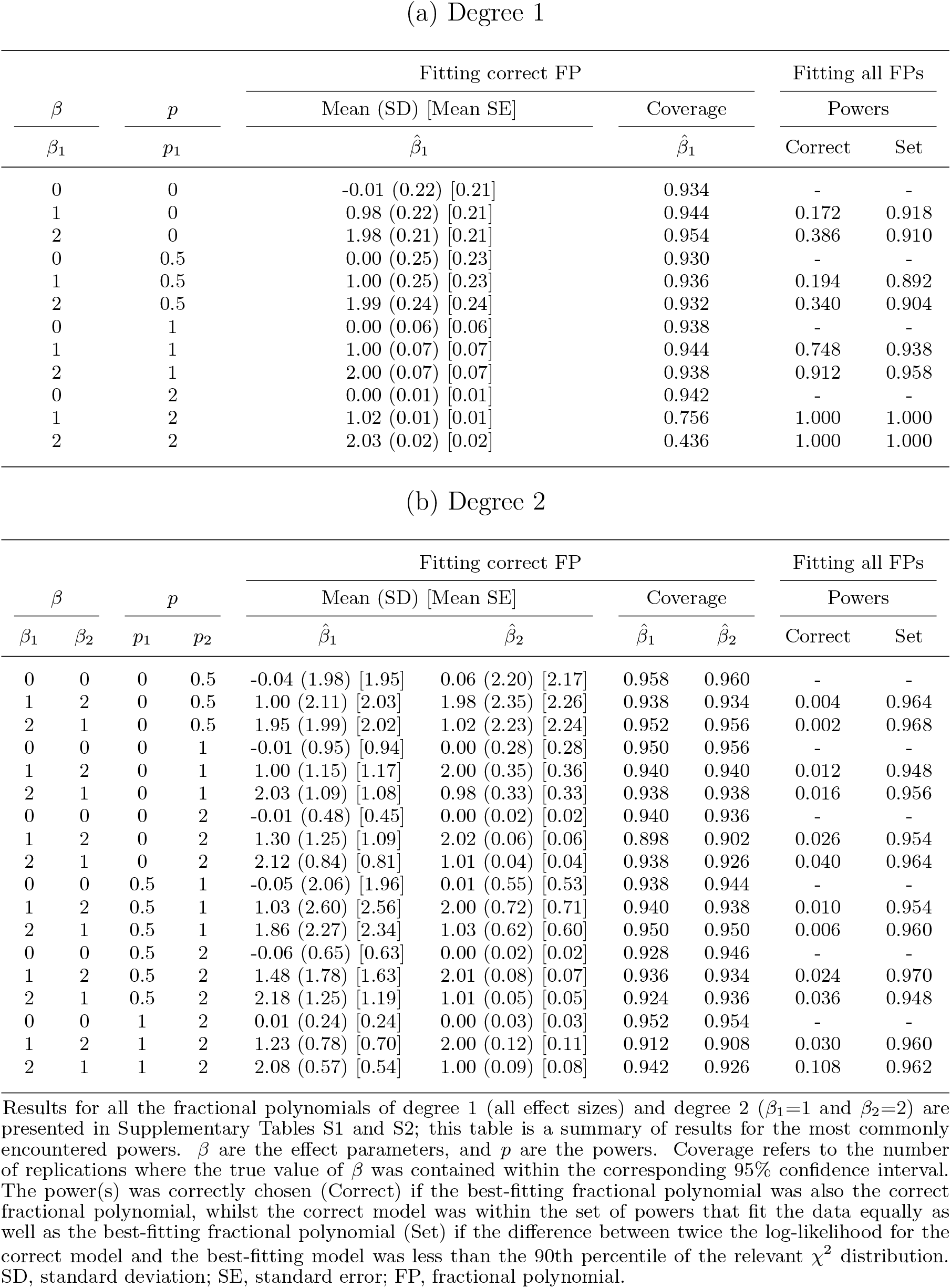
Simulation results for the fractional polynomial method.

For fractional polynomials of degree 1, when fitting the correct fractional polynomial model, the causal estimate was generally unbiased (Table 1). Coverage estimates were close to the nominal 95% rate, except for fractional polynomials of power 2 (and power 3, Supplementary Table S1), where causal estimates were slightly biased, and this small bias led to undercoverage. However, under the null, causal estimates were unbiased and correct coverage rates were maintained. For fractional polynomials of degree 2, a similar pattern was observed except that small biases and resulting undercoverage was more common, although correct coverage rate under the null was always maintained.

When fitting all the fractional polynomial models, the correct fractional polynomial model was fitted more often for a fractional polynomial of degree 1, and when the power of the fractional polynomial differed substantially from 0. However, in all cases, the correct fractional polynomial was in the set of best-fitting fractional polynomials in at least 89% of simulations. Under the null, the probability of fitting the ‘correct’ fractional polynomial was not estimated as all fractional polynomials with zero coefficients would describe the data equally well.

### Piecewise linear method

The piecewise linear method performed well when the true model was piecewise linear (such as a linear or a threshold relationship), with the predicted mean values of the outcome similar to their true values at the mean value of the exposure within each decile of the IV-free exposure (Table 2). The bootstrapped confidence intervals also had approximately 95% coverage at these points, except for the quantiles at or either side of the point of inflection of the threshold model. However, when the true model was not piecewise linear (in particular, for a quadratic relationship), estimates were biased and coverage was below nominal levels.

**Table 2.**
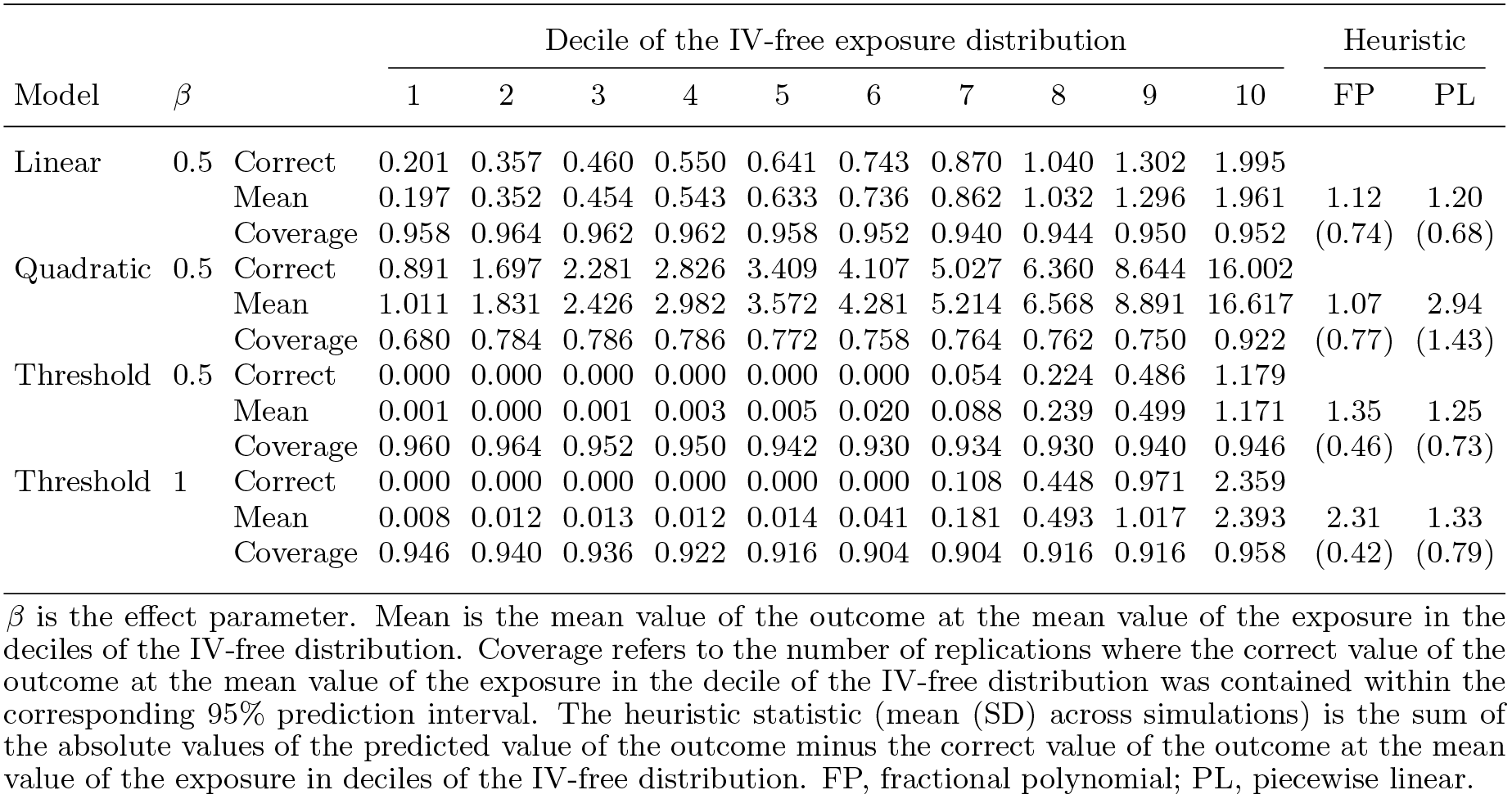
Simulation results for the piecewise linear method.

| Model | $\beta$ | | Decile of the IV-free exposure distribution | | | | | | | | | | Heuristic | |
| --- | --- | --- | --- | --- | --- | --- | --- | --- | --- | --- | --- | --- | --- | --- |
|  |  |  | 1 | 2 | 3 | 4 | 5 | 6 | 7 | 8 | 9 | 10 | FP | PL |
| Linear | 0.5 | Correct | 0.201 | 0.357 | 0.460 | 0.550 | 0.641 | 0.743 | 0.870 | 1.040 | 1.302 | 1.995 |  |  |
|  |  | Mean | 0.197 | 0.352 | 0.454 | 0.543 | 0.633 | 0.736 | 0.862 | 1.032 | 1.296 | 1.961 | 1.12 | 1.20 |
|  |  | Coverage | 0.958 | 0.964 | 0.962 | 0.962 | 0.958 | 0.952 | 0.940 | 0.944 | 0.950 | 0.952 | (0.74) | (0.68) |
| Quadratic | 0.5 | Correct | 0.891 | 1.697 | 2.281 | 2.826 | 3.409 | 4.107 | 5.027 | 6.360 | 8.644 | 16.002 |  |  |
|  |  | Mean | 1.011 | 1.831 | 2.426 | 2.982 | 3.572 | 4.281 | 5.214 | 6.568 | 8.891 | 16.617 | 1.07 | 2.94 |
|  |  | Coverage | 0.680 | 0.784 | 0.786 | 0.786 | 0.772 | 0.758 | 0.764 | 0.762 | 0.750 | 0.922 | (0.77) | (1.43) |
| Threshold | 0.5 | Correct | 0.000 | 0.000 | 0.000 | 0.000 | 0.000 | 0.000 | 0.054 | 0.224 | 0.486 | 1.179 |  |  |
|  |  | Mean | 0.001 | 0.000 | 0.001 | 0.003 | 0.005 | 0.020 | 0.088 | 0.239 | 0.499 | 1.171 | 1.35 | 1.25 |
|  |  | Coverage | 0.960 | 0.964 | 0.952 | 0.950 | 0.942 | 0.930 | 0.934 | 0.930 | 0.940 | 0.946 | (0.46) | (0.73) |
| Threshold | 1 | Correct | 0.000 | 0.000 | 0.000 | 0.000 | 0.000 | 0.000 | 0.108 | 0.448 | 0.971 | 2.359 |  |  |
|  |  | Mean | 0.008 | 0.012 | 0.013 | 0.012 | 0.014 | 0.041 | 0.181 | 0.493 | 1.017 | 2.393 | 2.31 | 1.33 |
|  |  | Coverage | 0.946 | 0.940 | 0.936 | 0.922 | 0.916 | 0.904 | 0.904 | 0.916 | 0.916 | 0.958 | (0.42) | (0.79) |

$\beta$ is the effect parameter. Mean is the mean value of the outcome at the mean value of the exposure in the deciles of the IV-free distribution. Coverage refers to the number of replications where the correct value of the outcome at the mean value of the exposure in the decile of the IV-free distribution was contained within the corresponding 95% prediction interval. The heuristic statistic (mean (SD) across simulations) is the sum of the absolute values of the predicted value of the outcome minus the correct value of the outcome at the mean value of the exposure in deciles of the IV-free distribution. FP, fractional polynomial; PL, piecewise linear.

Using the heuristic function (3) to compare between the estimates for the best-fitting fractional polynomial and the piecewise linear model, the models performed similarly under a linear model. For a quadratic model, the fractional polynomial method out-performed the piecewise linear method, whereas the opposite was true for a threshold model. This is unsurprising, as the fractional polynomial method performed best when the true model was a polynomial and likewise for the piecewise linear method when the model was piecewise linear.

### Varying the number of strata

The best-fitting fractional polynomial method had a similar or slightly better model fit (judged by the heuristic function) when a greater number of strata were used (Table 3). However, the piecewise linear method fitted the data better when fewer strata were used. Whereas the fractional polynomial method ensures that the estimate of the exposure-outcome relationship is a smooth function regardless of the number of strata, the estimate from the piecewise linear method becomes increasingly jagged as the number of strata increases.

**Table 3.**
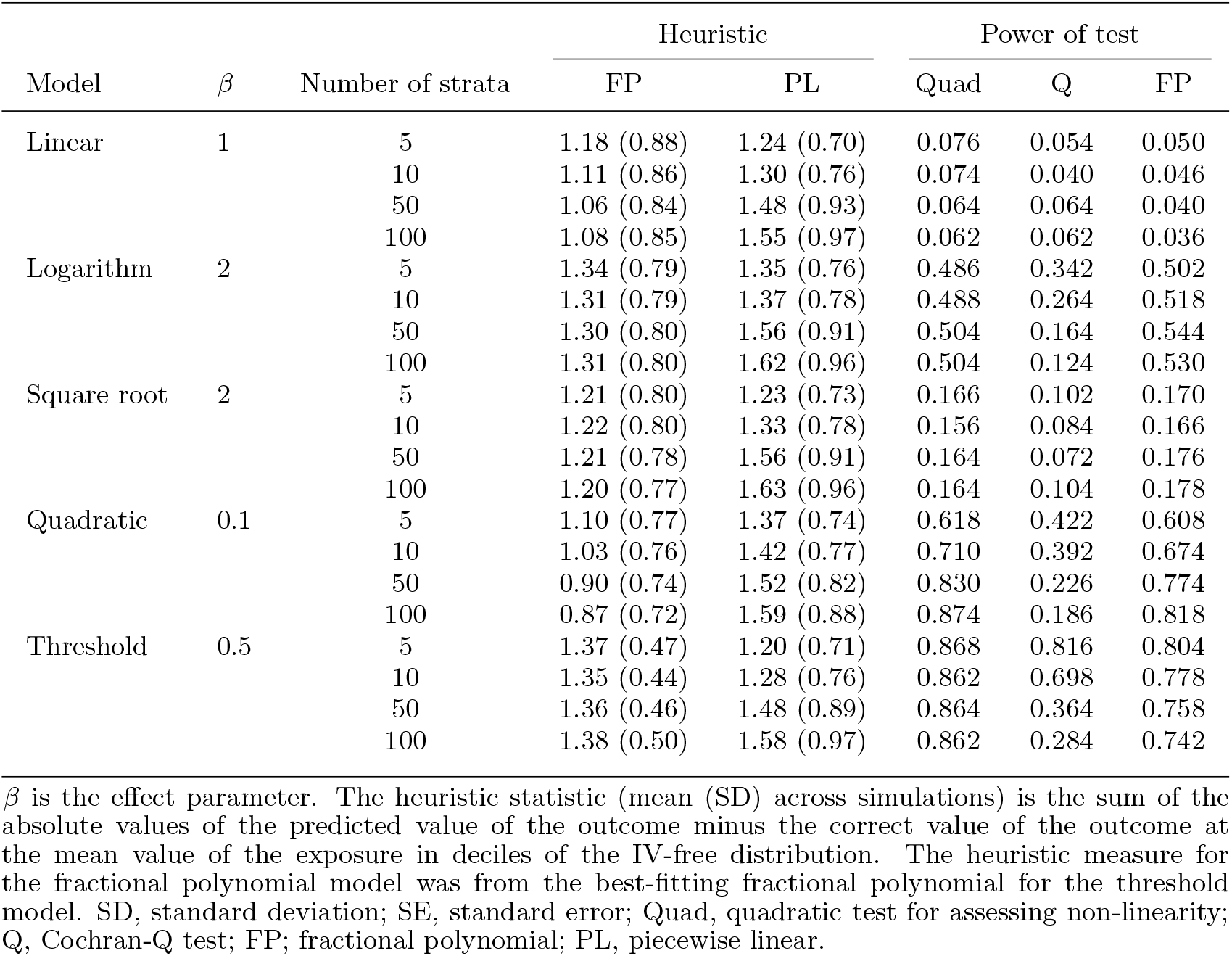
Simulation results for the piecewise linear method.

The coverage under the null (i.e. a linear model) was not overly inflated for any of the tests. In general, the fractional polynomial and quadratic tests were more powerful than the Cochran Q test across the simulations. The power of the Cochran Q test also decreased as the number of strata increased, whereas the power of the other tests either remained the same or increased. The quadratic test slightly outperformed the fractional polynomial test when the true model was a quadratic or a threshold model; the fractional polynomial test was slightly superior when the true model was a logarithm or a square root model.

### Additional simulations to assess impact of violations of assumptions

In the simulations where we relaxed the assumptions that the IV-exposure and the exposure-outcome effects are the same for all individuals, we found that the fractional polynomial models of degree 1 and the piecewise linear method both performed well in terms of bias and coverage (Supplementary Tables S3 and S4). The only concern was that tests of non-linearity had slightly inflated Type I error rates when the IV-exposure and exposure-outcome effects were varied in a correlated way; Type I error rate inflation was not observed when the effects were varied either separately or independently.

## Application of methods to the relationship between body mass index and blood pressure in UK Biobank

We illustrate the methods proposed in this paper in an applied example considering the shape of relationship between BMI and blood pressure in the UK Biobank study. UK Biobank is a prospective cohort study of 502,682 participants recruited at 22 assessment centres across the UK between 2006 and 2010 [Sudlow et al., 2015]. Participants were aged between 40 and 69 at baseline. Extensive health, lifestyle, biological and genetic measurements were taken on all participants. At the time of writing this paper, genetic information was only available for 133,687 individuals of European ancestry. For individuals on anti-hypertensive medication, 15/10mmHg were added to their SBP/DBP measurement, respectively. A sensitivity analysis was performed in individuals who had no history of hypertension.

To create an allele score (also called a genetic risk score) of variants related to BMI to be used as an instrumental variable, we extracted the 97 variants previously associated with BMI at a genome-wide level of significance by the GIANT consortium [Locke et al., 2015]. A proxy variant (rs751414; *r*^2^ = 0:99) was used instead of rs2033529, as this variant was not available in UK Biobank; the linkage disequilibrium information was calculated using the European samples from 1000 Genomes [1000 Genomes Project Consortium et al., 2012]. All of the variants were either directly genotyped or well-imputed (INFO>0.9). The allele score for each individual was computed by multiplying the number of BMI-increasing alleles for each variant by the effect of the variant on BMI (as estimated in the GIANT consortium) and summing across the 97 variants [Burgess and Thompson, 2013]. Overall this score explained 1.75% of the variance in BMI. We performed both the fractional polynomial and piecewise linear methods for estimating the relationships of BMI with systolic blood pressure (SBP) and diastolic blood pressure (DBP). The fractional polynomial method was implemented using 100 strata, whereas the piecewise linear method was implemented using 10 strata to avoid the exposure-outcome curve being overly jagged. The reference point was set at 25kg/m^2^.

To account for the multiple centres, we standardized the measure of BMI by stratifying individuals based on their residual value of BMI (the IV-free exposure) after regression of BMI on the allele score, age, sex, and centre. Adjustment for age, sex and centre was also made in the regressions to obtain the LACE estimates in each quantile group.

To assess the assumption that the effect of the IV on BMI is constant over the entire distribution of BMI, we also considered BMI as the outcome and calculated the associations of the IV with BMI in each of the strata. We then conducted tests (trend and Cochran Q tests) to investigate heterogeneity in the IV associations with BMI in different strata.

### Results of applied example

The exposure-outcome relationships for BMI with SBP and DBP estimated using the fractional polynomial and piecewise linear methods are presented in Figure 1. There were strong causal effects of BMI on both SBP and DBP (*p*-value < 1 × 10^−5^ for the causal estimates differing from zero in the fractional polynomial methods). There was strong evidence that the association between BMI and SBP was non-linear, with the quadratic test yielding *p*-value = 0.0026 (*p*-value = 0.0164 for the fractional polynomial test, *p*-value = 0.0346 for the Cochran Q test). The best-fitting fractional polynomial of degree 1 for the relationship between BMI and SBP had power −0.5, and there was no evidence to suggest that a fractional polynomial of degree 2 fitted the data better (*p*-value = 0.135). The estimate of the exposure-outcome relationship from the piecewise linear method visually suggested a threshold-type relationship, with a steep slope up to a BMI value of about 28kg/m^2^, and a slightly negative slope from 28kg/m^2^ onwards. The relationship between BMI and SBP was similar in individuals with no history of hypertension (Supplementary Figure S1).

**Figure 1 Figure 1:**
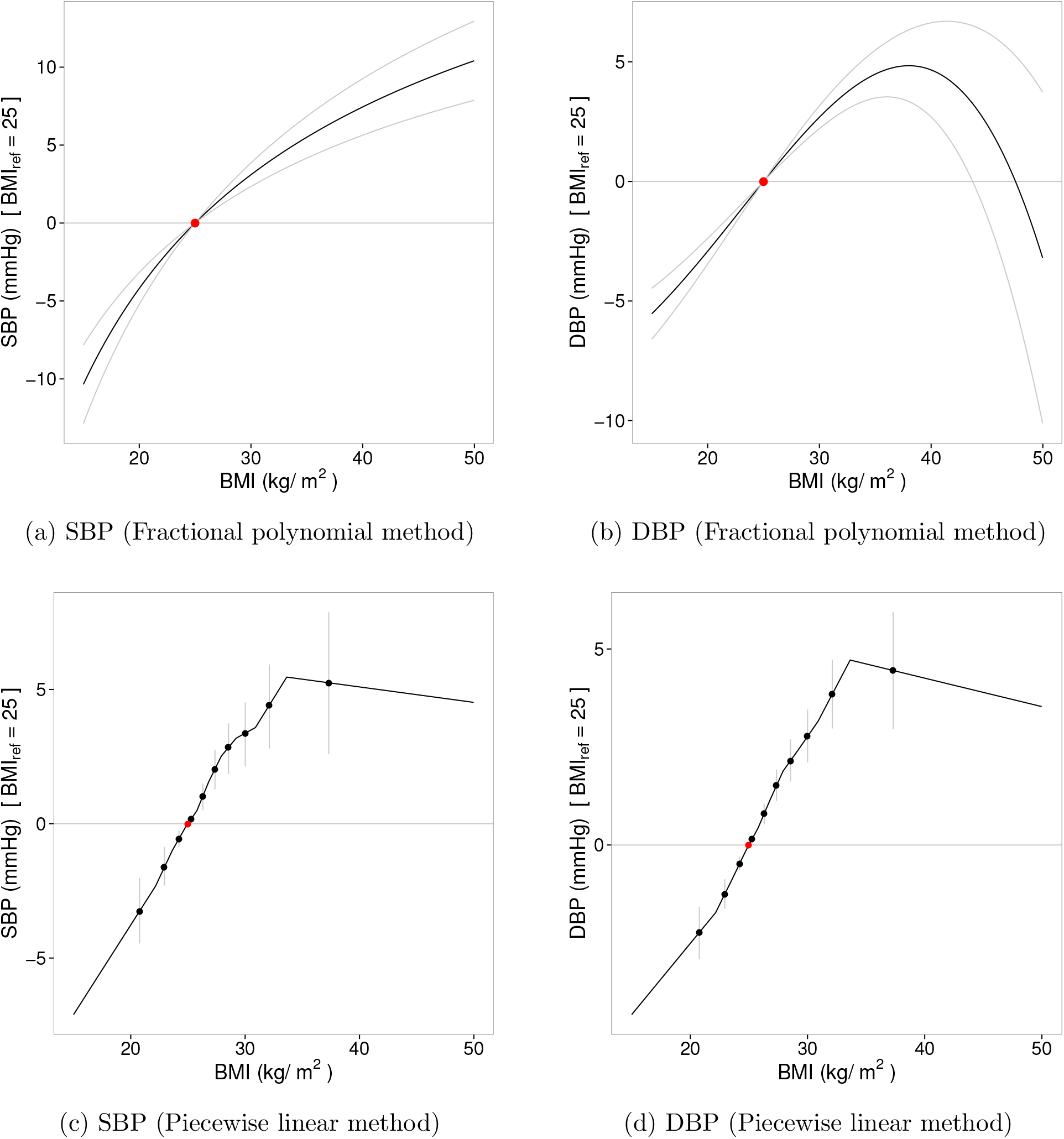
Causal effects of body mass index (BMI) on blood pressure (systolic blood pressure, SBP; diastolic blood pressure, DBP) using the fractional polynomial and piecewise linear methods on data from UK Biobank. The red point represents the reference point of BMI of 25 kg/m^2^. Grey lines represent 95% CIs. The fractional polynomial method used 100 strata.

The association between BMI and DBP was also non-linear (quadratic test *p*-value = 0.0005, fractional polynomial test *p*-value = 0.0114, Cochran Q test *p*-value = 0.0049), and there was strong evidence that the best-fitting fractional polynomial of degree 2 (with *p*_1_ and *p*_2_ = 3) fitted the data better than the best-fitting fractional polynomial of degree 1 (*p*-value = 0.0062). There was no evidence of a different relationship between BMI and DBP for underweight individuals, with the exposure-outcome curve increasing almost linearly up to a BMI of around 40kg/m^2^. But for hyper-obese individuals (BMI > 40kg/m^2^), DBP seemed to decrease sharply. This was particularly evident in the fractional polynomial method, which used a greater number of strata and hence had more resolution to consider the shape of the exposure-outcome relationship at the extremes of the BMI distribution. One potential reason for this finding is that hyper-obese individuals with high DBP are less likely to be enrolled in UK Biobank, perhaps due to differential survival probability. Another reason could be the difficulties in estimating blood pressure in hyper-obese individuals [Leblanc et al., 2013]. However, there was no evidence that the relationship between BMI and DBP was non-linear in individuals with no history of hypertension (*p*-value > 0.05 for all tests; Supplementary Figure S1).

There was no evidence that the associations of the IV with BMI varied between the different strata (trend test *p*-value = 0.135, Cochran Q test *p*-value = 0.901).

## Discussion

In this paper, we have proposed and tested two novel methods for examining the relationship between an exposure and an outcome using instrumental variable analysis in the context of Mendelian randomization. Both methods rely on stratifying the population based on the IV-free exposure; the exposure minus the effect of the IV. A causal effect, referred to as a LACE, is estimated in each stratum of population. The first method performs meta-regression on these LACE estimates using fractional polynomials. The second method estimates a continuous piecewise linear function, the gradient of which in each stratum is the LACE estimate for that stratum. Both methods were demonstrated in a simulation study to estimate the true exposure-outcome relationship well when its functional form corresponded to the form of the estimate from each method (that is, when the exposure-outcome relationship was a fractional polynomial for the fractional polynomial method, and when the relationship was piecewise linear for the piecewise linear method), with causal estimates being close to unbiased and coverage rates generally maintaining nominal levels (in particular, coverage rates were always correct under the null). Additionally, tests of non-linearity were provided and their performance was assessed. The quadratic and fractional polynomial tests had the best performance in terms of Type I error rate and power.

The recommendation as to which method to use depends on the aim of the investigation. The fractional polynomial method will always provide a smooth estimate of the exposure-outcome relationship, and as such has more consistent performance when a large number of strata are chosen (i.e. when the shape of the relationship is considered over a wider and more detailed range of the exposure distribution). Fractional polynomials of degree 1 had better performance than those of degree 2 in terms of bias and coverage of effect estimates. However, fractional polynomials of degree 1 are less flexible and would not be able to model complex exposure-outcome relationships. Additionally, they tend to smooth over discrepancies in the data. For example, if the LACE estimate for individuals in the lowest quantile group for BMI was substantially different to the other LACE estimates, then both this difference and any uncertainty in the LACE estimate would be smoothed over somewhat in the fractional polynomial estimate. Preference between the methods therefore comes down to a question of prior belief: if one truly believes the true exposure-outcome relationship to be smooth, and that estimates in the surrounding quantiles should be used to model the LACE in the target quantile, then the fractional polynomial method should be preferred. However, if one does not want to smooth over estimates, then the piecewise linear method should be preferred; however, the estimate of the exposure-outcome relationship will be more jagged and variable.

### Interpretation of the exposure-outcome relationship

If the function relating the exposure to the average value of the outcome is homogeneous across the population, then the methods provided in this paper estimate this function (the exposure-outcome relationship) even if there is unmeasured confounding. If the function is heterogeneous, then the situation is more complicated [Small, 2014]. For example, taking BMI as the exposure, if the subject-specific effect curve (as defined by Small) is linear for all individuals in the population, but the magnitude of effect is greater for overweight individuals, then the exposure-outcome relationship will be quadratic (or at least convex and positive) rather than linear. The exposure-outcome curve at low values of the exposure is only estimated using underweight individuals, and at high values of the exposure only using overweight individuals. However, this is perhaps the most relevant way to express the exposure-outcome relationship, as the causal effect of reducing one’s BMI from 20kg/m^2^ to 18kg/m^2^ is not so relevant for someone with a BMI of 40kg/m^2^. Hence, we do not claim any global interpretation of the exposure-outcome relationship as estimated in this paper apart from in the unlikely case that the functional relationship is homogeneous in the population. It is better interpreted as a series of local estimates, which are graphically connected in order to compare and contrast trends in these local estimates at different values of the exposure, and to compare the relative benefit of intervening on the exposure for individuals with different values of the exposure, but which does not necessarily reflect the effect of intervening on the exposure to take any value in its distribution for any single individual.

### Measurement error in the exposure

As has been noted in other contexts, estimates of non-linear relationships are sensitive to measurement error in the exposure [Keogh et al., 2012]. The standard ‘triple whammy’ of measurement error is likely to apply here: measurement error biases parameter estimates, reduces power, and obscures important features in the shape of relationships [Carroll et al., 2006]. For example, with a threshold relationship, measurement error in the exposure would mean that the point of inflexion in the exposure-outcome relationship would be less sharply evident. In the case of BMI, measurement error is not such an issue, as height and weight can be measured precisely, and neither variable experiences substantial diurnal or seasonal variation. However, for other exposures, measurement error may affect results more severely.

### Requirement of concomitant and individual-level data

Many recent advances in Mendelian randomization have enabled investigations to be performed using summarized data on the genetic associations with the exposure and with the outcome only, and/or in a two-sample setting in which genetic associations with the exposure and with the outcome are estimated on separate groups of individuals [Burgess et al., 2013, 2015]. However, estimation of the exposure-outcome relationship requires both individual-level data and a one-sample setting (otherwise neither stratification of the population nor the estimation of genetic associations with the outcome in the strata are possible). Although, large cohorts with concomitant data on genetic variants, exposures, and outcomes are becoming more widely available, particularly in the form of biobanks such as UK Biobank.

In conclusion, these two novel methods are useful in investigating non-linear exposure-outcome relationships. The methods allow for easy graphical assessment of the shape of the relationship, and allied with tests of non-linearity, provide an effective tool for assessing non-linear exposure-outcome relationships using IV analysis for Mendelian randomization.

## Acknowledgements

This research has been conducted using the UK Biobank Resource (Application 7439).

This work was supported by the UK Medical Research Council [G66840, G0800270], Pfizer [G73632], British Heart Foundation [SP/09/002], UK National Institute for Health Research Cambridge Biomedical Research Centre, European Research Council [268834], and European Commission Framework Programme 7 [HEALTH-F2-2012-279233]. Stephen Burgess is supported by the Wellcome Trust [100114].

